# Transient Reprogramming of Neonatal Cardiomyocytes to a Proliferative Dedifferentiated State

**DOI:** 10.1101/801092

**Authors:** Thomas Kisby, Irene de Lázaro, Maria Stylianou, Giulio Cossu, Kostas Kostarelos

**Affiliations:** Nanomedicine Lab, Faculty of Biology, Medicine and Health, AV Hill Building, The University of Manchester, Manchester, M13 9PT, UK; Division of Cell Matrix Biology and Regenerative Medicine, Faculty of Biology, Medicine and Health, The University of Manchester, Manchester, M13 9PT, UK; Harvard John A. Paulson School of Engineering and Applied Sciences, 58 Oxford Street, Cambridge, MA 02138, USA; Wyss Institute for Biologically Inspired Engineering at Harvard University, Center for Life Science, Boston, MA 02115, USA

**Keywords:** Cell reprogramming, Cardiomyocytes, OSKM, Pluripotency, Adenovirus, Gene therapy

## Abstract

Zebrafish and urodele amphibians are capable of extraordinary myocardial regeneration thanks to the ability of their cardiomyocytes to undergo transient dedifferentiation and proliferation. Somatic cells can be temporarily reprogrammed to a proliferative, dedifferentiated state through transient expression of *Oct3/4, Sox2, Klf4* and *c-Myc* (OSKM) transcription factors. Here, we utilized an OSKM-encoding non-integrating vector to induce transient reprogramming of mammalian cardiomyocytes *in vitro*. Reprogramming factor expression in neonatal rat cardiomyocytes triggered rapid cell dedifferentiation characterized by downregulation of cardiomyocyte specific gene and protein expression, sarcomere dis-assembly and loss of autorhythmic contractile activity. Concomitantly, a significant increase in cell cycle related gene expression and Ki67 positive cells was observed, indicating that dedifferentiated cardiomyocytes possess an enhanced proliferative capacity. A small proportion of cardiomyocytes progressed through mesenchymal to epithelial transition, further indicating the initiation of cell reprogramming. However, complete reprogramming to a pluripotent-like state was not achieved for the duration of the study (20 days), both in standard and embryonic stem cell culture media conditions. The transient nature of this partial reprogramming response was confirmed as cardiomyocyte-specific cell morphology, gene expression and contractile activity were recovered by day 15 after viral transduction. Further investigations into the complete downstream biological effects of ectopic OSKM expression in cardiomyocytes and the fate of these reprogrammed cells are warranted. Our results to date suggest that transient reprogramming could be a feasible strategy to recapitulate regenerative mechanisms of lower vertebrates and inform direct gene therapy approaches to cardiac regenerative medicine.

## Introduction

The majority of cardiomyocytes in the mammalian heart exit the cell cycle soon after birth (Soonpaa et al., 1996, Walsh et al., 2010). Although evidence exists for the renewal of a small percentage of cardiomyocytes throughout the lifetime of mammals (Bergmann et al., 2009, Senyo et al., 2013, Bergmann et al., 2015), such low rates cannot effectively support myocardial regeneration following an ischemic insult. In contrast, zebrafish have a much greater capacity for cardiomyocyte replenishment following various forms of cardiac injury (Poss et al., 2002, Raya et al., 2003, González-Rosa et al., 2011). Lineage tracing studies in these organisms have confirmed that pre-existing cardiomyocytes are the primary source of regenerated myocardium, indicating that extensive cardiomyocyte proliferation takes place after injury (Kikuchi et al., 2010). Additionally, it has been observed that cardiomyocyte cell cycle re-entry coincides with partial dedifferentiation to a progenitor-like state (Jopling et al., 2010, Sleep et al., 2010, Sallin et al., 2015, Wu et al., 2016). These observations have also been made in urodele amphibians which possess a comparable regenerative advantage (Laube et al., 2006). A similar albeit more subtle response is observed in embryonic and neonatal mammals however, this regenerative capacity is quickly lost within 7 days after birth (Porrello et al., 2011, Drenckhahn et al., 2008, Lam and Sadek, 2018). Inducing partial dedifferentiation of mammalian cardiomyocytes could therefore be an attractive strategy to enhance the proliferative capacity of these cells.

A defined combination of transcription factors; *Oct3/4, Sox2, Klf4* and *cMyc* (OSKM), is thought to be universal in its ability to reset the epigenetic modulations that maintain cells in their differentiated state (Takahashi and Yamanaka, 2006). Indeed, many cell types including cardiomyocytes have been completely reprogrammed to an embryonic stem cell (ESC)-like state, known as induced pluripotent stem cells (iPSCs), through the forced expression of these factors (Rizzi et al., 2012, Xu et al., 2012, Cheng et al., 2017). To reach *bona fide* iPSCs, cells must first progress through various phases of reprogramming defined as initiation, maturation and stabilisation (Samavarchi-Tehrani et al., 2010, Tanabe et al., 2013). In order to advance through these stages and eventually generate stable, transgene independent iPSCs, OSKM overexpression needs to be maintained for at least 8-12 days (Brambrink et al., 2008, Samavarchi-Tehrani et al., 2010). If OSKM expression is withdrawn earlier, cells reach instead a state of partial or incomplete reprogramming that retain transcriptional and epigenetic marks from the cell type of origin (Guo et al., 2017, Polo et al., 2012, Stadtfeld et al., 2008a). The spontaneous re-differentiation of partially reprogrammed cells to their cell type of origin following OSKM withdrawal has been frequently reported (Brambrink et al., 2008, Woltjen et al., 2009, Samavarchi-Tehrani et al., 2010, Tanabe et al., 2013). For such reasons, partially reprogrammed cells are increasingly recognised as potential novel sources of lineage restricted progenitor-like cells for cell therapy applications (Guo et al., 2017, Zhang et al., 2016).

While partial and transient OSKM mediated reprogramming of several cell types has been demonstrated, the response of cardiomyocytes to short term OSKM expression is yet to be explored. Here, we aimed to understand whether cardiomyocytes could be partially reprogrammed *in vitro* using adenoviral mediated OSKM expression. We further investigated the proliferative capacity of reprogrammed cardiomyocytes and explored their propensity to return to their original phenotype in the absence of additional differentiation stimuli.

## Results

### Efficient Expression of OSKM in Neonatal Rat Cardiomyocytes using a Polycistronic Adenoviral Vector

To investigate the response of cardiomyocytes to OSKM expression we utilized primary neonatal rat cardiomyocytes (NRCMs) extracted from post-natal day 2 Sprague Dawley rats. The purity of these cultures at the time of transduction was >80% as determined by cardiac troponin-T (cTnT) and NKX2-5 expression (**Figure S1a-c**), which is comparable with other reports (Mohamed et al., 2018). To force the expression of OSKM in these cells and interrogate partial reprogramming, we used an adenoviral vector (Ad-CMV-MKOS) containing all four reprogramming genes in a single polycistronic expression cassette (**Figure 1a**). We hypothesised that, based on their non-integrating nature, adenoviral vectors would provide transient OSKM expression compatible with transient cell reprogramming, as opposed to the sustained expression achieved with integrating viral vectors. Indeed, previous reports using adenoviral vectors for cellular reprogramming evidenced the need for repeat transduction to sustain OSKM gene expression long enough to generate iPSCs (Stadtfeld et al., 2008b, Zhou and Freed, 2009, Okita et al., 2008). We first investigated mRNA expression of total *Oct3/4, Sox2, Klf4* and *cMyc* 3 days post transduction with Ad-CMV-MKOS (**Figure 1b**). All four genes were overexpressed in NRCMs by relatively low multiplicities of infection (MOIs), consistent with reports of the high amenability of this cell type to adenoviral mediated gene delivery (Mohamed et al., 2018, Louch et al., 2011). Protein expression of OCT3/4 and SOX2 was also confirmed by immunocytochemistry at this same timepoint (**Figure 1c**). This was used to determine transduction efficiency (circa 37%) based on the number of SOX2 positive nuclei at the 5 PFU/cell dose (**Figure 1d**). An empty adenoviral vector (Ad-CMV-Null) was used throughout our study to ensure that changes observed were due to OSKM expression and not due to viral infection alone. Indeed, we did not observe the presence of OCT3/4 or SOX2 positive cells in Ad-CMV-Null treated cultures (**Figure 1c**). We next followed the expression of reprogramming factors over time. Adenoviral mediated *OSKM* expression peaked 3-5 days post transduction however, this level of expression was not sustained (**Figure 1e**). By day 10, the levels of *OKSM* mRNA decreased toward baseline consistent with a non-integrating vector. Similar expression kinetics were observed at the protein level with the highest number of SOX2 positive cells observed 3 days after transduction (**Figure 1f-g**). Therefore, we focused on these timepoints to understand the early responses of cardiomyocytes to OSKM overexpression.

**Figure 1:**
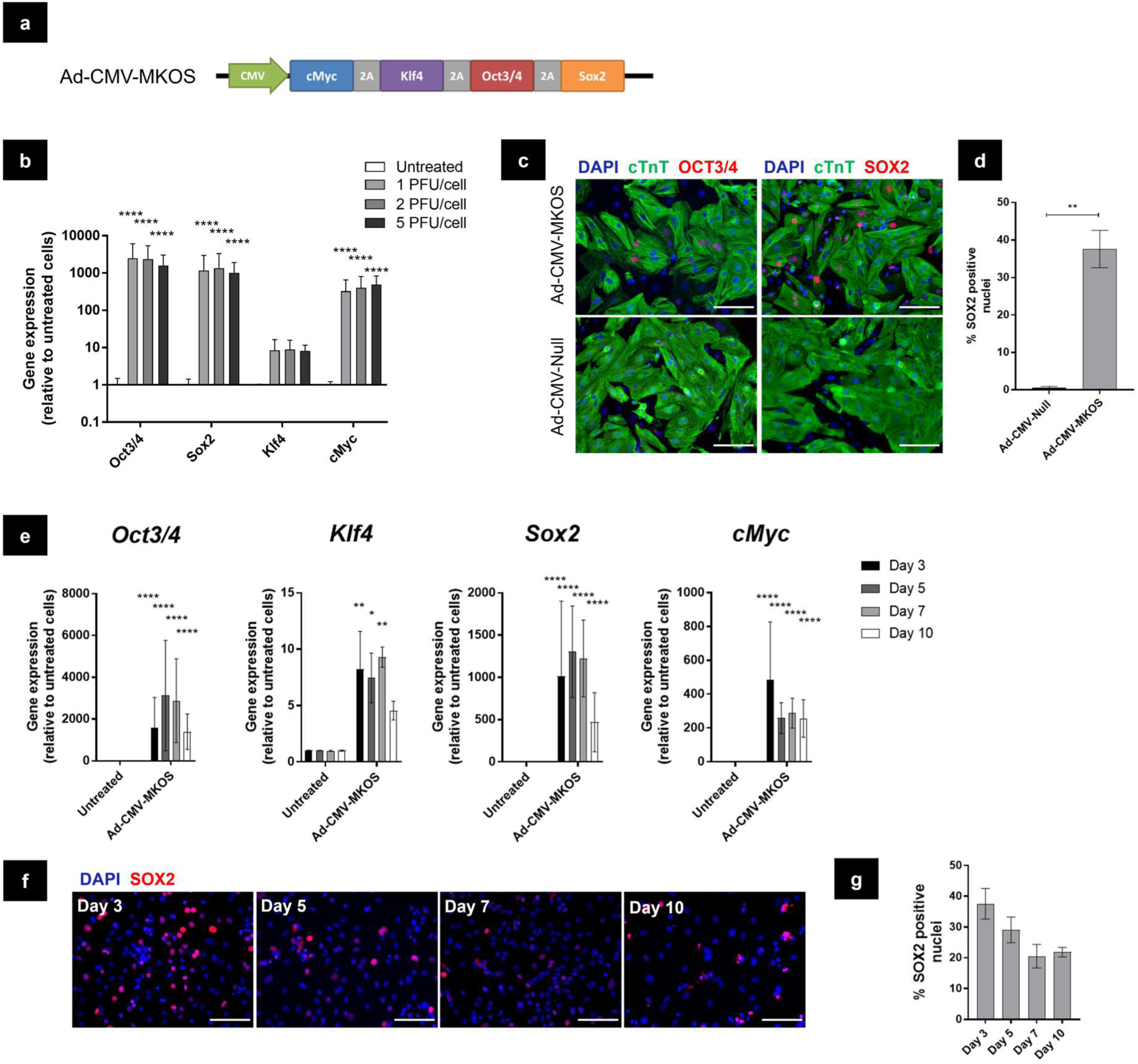
OSKM is Efficiently Delivered to Neonatal Rat Cardiomyocytes using a Polycistronic Adenoviral Vector. (**a**) Schematic of Ad-CMV-MKOS expression cassette. (**b**) Gene expression of total *OSKM* relative to untreated cells 3 days post transduction (n=3). (**c**) Representative images of OCT3/4 and SOX2 protein expression in cardiomyocytes stained with cTnT (scale bar = 100 µm). (**d**) Transduction efficiency expressed as percentage nuclei positive for SOX2 (n=3 replicates/4-6 fields per replicate). (**e**) Timecourse of total *OSKM* gene expression relative to untreated NRCMs (n=3). (**f**) Representative images of SOX2 positive cells on days 3 – 10 post transduction (scale bars = 100 µm). (**g**) Quantification of SOX2 positive nuclei (n=3 replicates, 4-6 fields per replicate). Data are presented as mean ± S.D. (**b**) and (**e**) one-way ANOVA with Tukey’s post hoc analysis, (**d**) Unpaired t-test, *, **, ***, and **** denote p < 0.05, p < 0.01, p < 0.001, and p < 0.0001, respectively.

### OSKM Overexpression Drives Rapid Dedifferentiation of Neonatal Cardiomyocytes

Three days post transduction (i.e coinciding with the peak levels of exogenous OSKM expression) we observed substantial morphological and functional changes in Ad-CMV-MKOS transduced NRCMs consistent with dedifferentiation. OSKM transduced NRCMs showed a flattened, less striated appearance with more prominent nuclei and nucleoli (**Figure 2a**). We also observed the cessation of autorhythmic contractile activity in the cells which possessed this altered morphology (**Video S1** and **Video S2**). Such morphological changes were further investigated by immunocytochemistry. Disassembly of sarcomeres within SOX2 positive cardiomyocytes was clearly observable however, this was largely absent from Ad-CMV-Null treated NRCMs (**Figure 2b**). In additional a visible reduction in cTnT expression was also apparent in a small number of OSKM transduced cardiomyocytes, indicating not only disassembly but also a reduction in the expression of sarcomere proteins (**Figure 2c**). Both phenomena recapitulate events characteristic of cardiomyocyte dedifferentiation in the zebrafish and newt regenerating myocardium (Jopling et al., 2010, Laube et al., 2006, Kikuchi et al., 2010). *Myh6* and *Myh7* are genes which encode α- and β-myosin heavy chain (MHC) respectively which are essential proteins involved in the contractile functions of adult and neonatal ventricular myocytes (Lompre et al., 1984). The expression of such transcripts was downregulated in OSKM transduced cultures (**Figure 2d**), but remained unaltered in NRCMs treated with the control vector confirming this response to be dependent on OSKM expression (**Figure S2a**). The downregulation of *Myh6* and *Myh7* occurred rapidly by day 3 post transduction but returned toward baseline levels concomitantly with the decrease in exogenous OSKM (**Figure 2d**). Additionally, a moderate upregulation of the pluripotency-associated genes *endogenous-Oct3/4* and *Nanog* was observed, although their expression levels were significantly lower than observed in a mouse ESC line and did not increase further overtime (**Figure 2e**). In agreement with these observations, the de-differentiation of NRCMs, characterised by sarcomere disorganization and loss of cTnT expression, appeared most prominent on days 3-5 post transduction (**Figure 2f**). By day 7 post transduction, NRCMs treated with Ad-CMV-MKOS showed more organised sarcomeres and homogenous cTnT expression (**Figure 2f***-***g**). Notably on day 10, the striated cardiomyocyte-like morphology was clear by phase contract microscopy (**Figure 2h**) and autorhythmic contractile activity was restored in these cells (**Videos S3** and **Video S4**). Overall, these data suggest that NRCMs can undergo OSKM-mediated dedifferentiation in line with the early stages of cell reprogramming.

**Figure 2:**
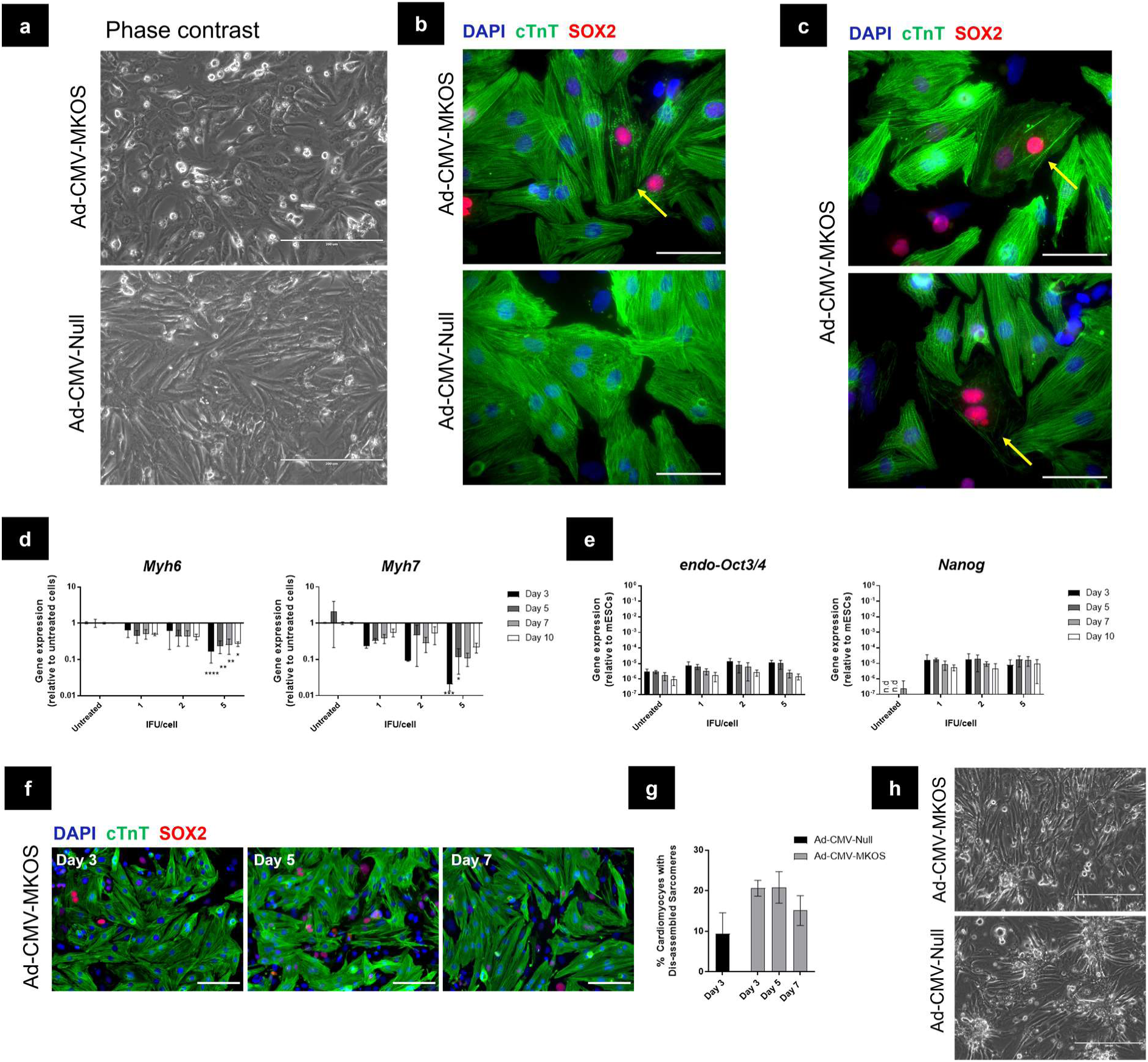
Overexpression of OSKM Induces Dedifferentiation of NRCMs. (**a**) Representative phase contrast image of NRCMs 3 days post transduction with Ad-CMV-MKOS or Ad-CMV-Null (scale bar = 200 µm) (n=5 replicates/ 3 fields per replicate). (**b**) Immunofluorescence of OSKM induced sarcomere disassembly (yellow arrow) (scale bar = 50 µm). (**c**) Reduction in cTnT expression in SOX2 positive cardiomyocytes (yellow arrows) (scale bar = 50 µm). (**d**) Gene expression of *Myh6* and *Myh7* after exposure to Ad-CMV-MKOS (n=3). (**e**) Gene expression of pluripotency related genes relative to mouse ESCs (n.d denotes transcript not detected) (n=3). (**f**) Immunofluorescence of NRCM morphology overtime following exposure to Ad-CMV-MKOS (scale bar = 100 µm). (**g**) Quantification of the percentage of cTnT positive cells showing disorganized sarcomere structure (n=3 replicates/4-6 fields per replicate). (**h**) Comparison between morphology of Ad-CMV-MKOS and Ad-CMV-Null treated cardiomyocytes 10 days post transduction (scale bar = 200 µm). Representative image from n=5 replicates, 3 fields per replicate. Data are presented as mean ± S.D. (**d**) and (**e**) one-way ANOVA with Tukey’s post hoc *, **, ***, and **** denote p < 0.05, p < 0.01, p < 0.001, and p < 0.0001, respectively.

To confirm the cardiomyocyte origin of OSKM transduced cells with reduced or absent cTnT expression, we utilized neonatal mouse cardiomyocytes (NMCMs) from an αMHC-Cre-tdTomato lineage tracing model. The expression of Cre recombinase under the transcriptional control of the cardiomyocyte-specific αMHC (*Myh6*) promoter permanently labels cardiomyocytes with the fluorescent protein tdTomato (Agah et al., 1997). Thus, OSKM transduced NMCMs can be unequivocally traced through their dedifferentiation. Following the same transduction protocol utilized for NRCMs, we observed sarcomere disassembly and reduced cTnT expression in tdTomato positive cells 3 days after transduction with Ad-CMV-MKOS (**Figure S3a**). The presence of SOX2+tdTomato+ cells with reduced cTnT expression confirm our previous observations that adenoviral mediated OSKM expression enables neonatal cardiomyocyte dedifferentiation. We did infrequently observe a small number of tdTomato+cTnT-cells in Ad-CMV-Null treated cultures however, these cells did not express SOX2 and may be attributed to Cre-recombinase activity in immature myocytes.

### Forced Dedifferentiation of Cardiomyocytes with OSKM Enhances Proliferative Capacity

It was next assessed whether OSKM induced dedifferentiation would enhance the capacity of NRCMs to undergo proliferation. We first assessed this using an alamarBlue cell viability assay that measures the metabolism mediated reduction of resazurin to a fluorescent product to estimate cell number and metabolic activity (Ahmed et al., 1994). OSKM expression increased the relative viability of NRCMs 3 days post transduction compared to cells treated with the control vector (**Figure 3a**) and this was maintained over time (**Figure 3b**). We used a haemocytometer to count cells following viability measurements which showed a similar trend indicating an increase in cell number in the OSKM transduced cultures (**Figure 3c**). Next, we investigated the expression of cell cycle related genes under the assumption that this would be altered if partially reprogrammed cardiomyocytes re-entered the cell cycle. Following OSKM overexpression, we observed an increase in *CyclinD1* (*Ccnd1*) expression concomitant with a downregulation of the negative cell cycle regulator *Cdkn2a* (**Figure 3d**). We next used Ki67 staining to assess the number of proliferating cells 3 days post transduction. We observed a significant increase in the total percentage of Ki67+ nuclei in NRCMs treated with Ad-CMV-MKOS compared to those treated with the control vector (**Figure 3e-f**). A smaller, but non-significant increase in the percentage of Ki67+cTnT+ cells also was observed (**Figure 3f**). Given our previous observation that some NRCMs lose cTnT expression following OSKM transduction, we concluded that this was not a reliable indicator of the number of dedifferentiated cardiomyocytes undergoing proliferation. Instead, we utilized αMHC-Cre-tdTomato NMCMs to confirm the source of proliferating cells and identified an increase in the percentage of Ki67+tdTomato+ cells. Such increase in percentage was consistent with that observed for total Ki67+ nuclei (**Figure S4a-b**). As expected, many Ki67+tdTomato+ cells showed reduced or absent cTnT expression (**Figure S4c**). Furthermore, in agreement with the hypothesis that OSKM mediated dedifferentiation drives cell cycle activation, co-localisation of Ki67 and OCT3/4 were observed within NRCMs that showed this dedifferentiated phenotype (**Figure 3g**). Overall, these data suggest that forced OSKM expression in NRCMs and NMCMs induces proliferation through cellular dedifferentiation.

**Figure 3:**
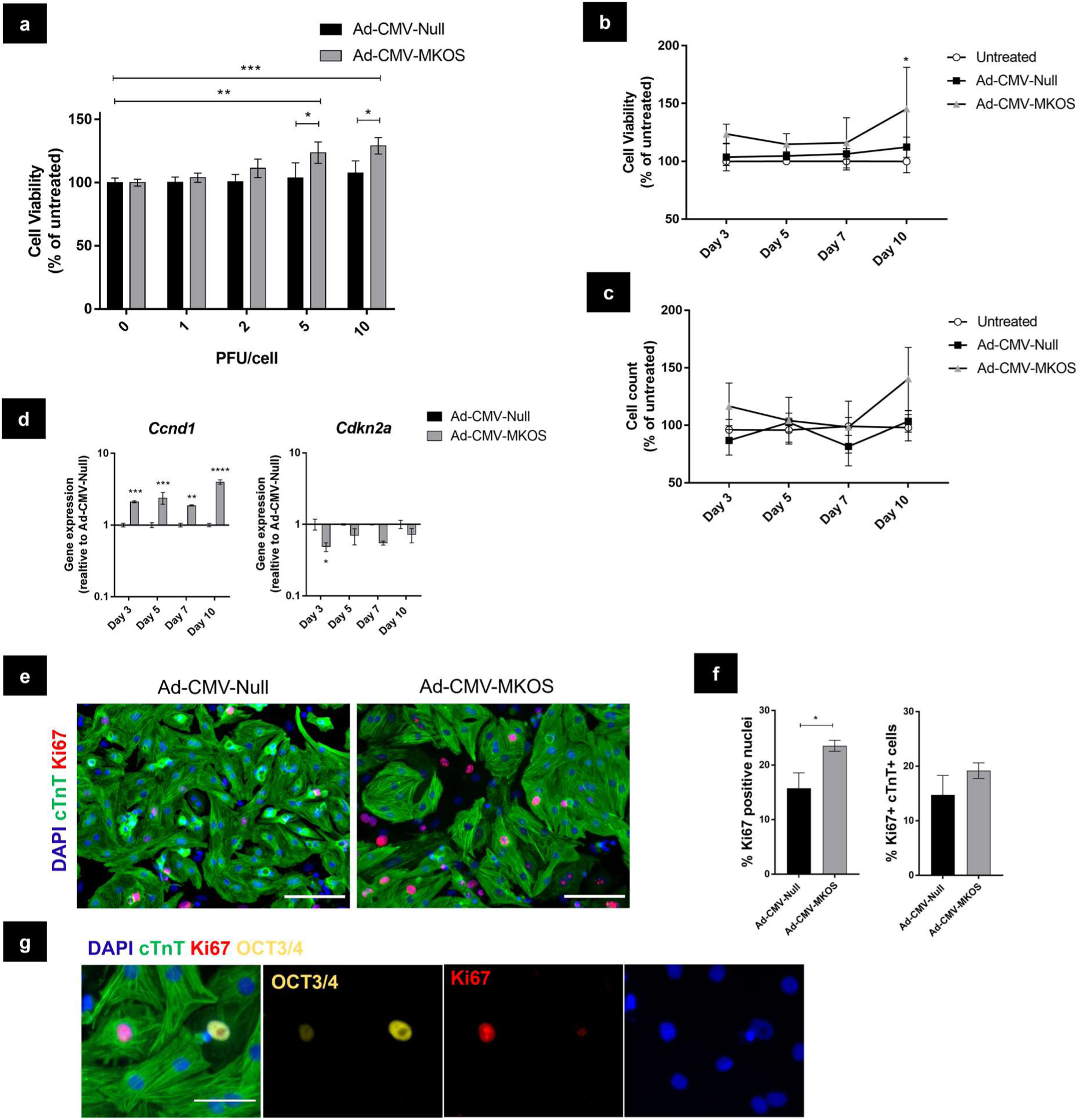
Forced Dedifferentiation of Cardiomyocytes by Reprogramming Enhances Proliferation. (**a**) Relative cell viability 3 days post transduction with Ad-CMV-Null or Ad-CMV-MKOS (n=3). (**b**) Timecourse of viability measurements at 5 PFU/cell (n=3). (**c**) Timecourse of cell counts at 5 PFU/cell (n=3). (**d**) Expression of cell cycle related genes (n=2). (**e**) Ki67 expression in NRCMs 3 days post transduction (Scale bar = 100 µm). (**f**) Quantification of total Ki67+ nuclei and cTnT+ Ki67+ cells (n=3 /4-6 fields per replicate). (**e**) Colocalisation of Ki67 within dedifferentiating OCT3/4+ cardiomyocytes (Scale bar = 50 um). Data are presented as mean ± S.D. (**a-d**) two-way ANOVA with Tukey’s post hoc analysis, (**f**) Unpaired t-test. *, ** and *** denote p < 0.05, p < 0.01 and p < 0.001 respectively.

### Cardiomyocytes are Partially but not Completely Reprogrammed by Adenoviral Mediated OSKM Expression

We next investigated the extent of cardiomyocyte reprogramming achieved by adenoviral mediated OSKM expression. In these experiments, we extended the timepoint of investigation up to 20 days post transduction, which is comparable to the length of conventional iPSC generation protocols (Takahashi and Yamanaka, 2006, Buganim et al., 2013). Mesenchymal to epithelial transition (MET) is a key event in the early stages of reprogramming (Samavarchi-Tehrani et al., 2010), during which stemness-associated makers including SSEA1 (a carbohydrate ESC marker that is a product of *Fut4*) are induced (Teshigawara et al., 2017). As early as 3 days after OSKM transduction, we observed an upregulation in epithelial associated genes *Cdh1* and *Epcam* in NRCMs treated with OSKM (**Figure 4a**). In contrast, no significant increases in the stem cell marker *Fut4* was detected at any of the timepoints investigated (**Figure 4a**). To confirm the increase in epithelial markers we stained cells for E-Cadherin (ECad) and identified a small number of ECad+ cells specifically in the OSKM treated cultures (**Figure 4b**). Consistent with *Cdh1* gene expression, the number of ECad+ cells did not increase over the timepoints investigated (**Figure 4c-d**). Notably the percentage of ECad+ cells was lower than the number of cells observed to be undergoing dedifferentiation (**Figure 4d**). Additionally, these cells did not appear to form cobblestone-like epithelial cell colonies suggesting a full transition to an epithelial phenotype had not taken place (**Figure 4c**). To determine the source of ECad+ cells, they were stained cells for the cardiac transcription factor NKX2-5 which is one of the earliest markers of the cardiac lineage (Schwartz and Olson, 1999). On day 3 post transduction, the majority of ECad+ cells expressed NKX2-5 suggesting cardiomyocytes were the primary source of these cells (**Figure 4e-f**). Furthermore, the maintenance of NKX2-5 expression in these cells further indicates that NRCMs do not pass completely through MET and still retain markers of their cardiomyocyte origin. To further confirm that Ad-CMV-MKOS transduction was able to induce partial MET in cardiomyocytes we used αMHC-Cre-tdTomato NMCMs. Consistent with the results for NRCMS, ECad+ cells were identified specifically in Ad-CMV-MKOS transduced cultures (**Figure S5a**). These ECad+ cells were also positive for tdTomato confirming that cardiomyocytes contribute to these partially reprogrammed cells (**Figures S5b**). Overall, these data suggest that a small number of cardiomyocytes initiate MET in line with the earliest stages of somatic cell reprogramming (Samavarchi-Tehrani et al., 2010).

**Figure 4:**
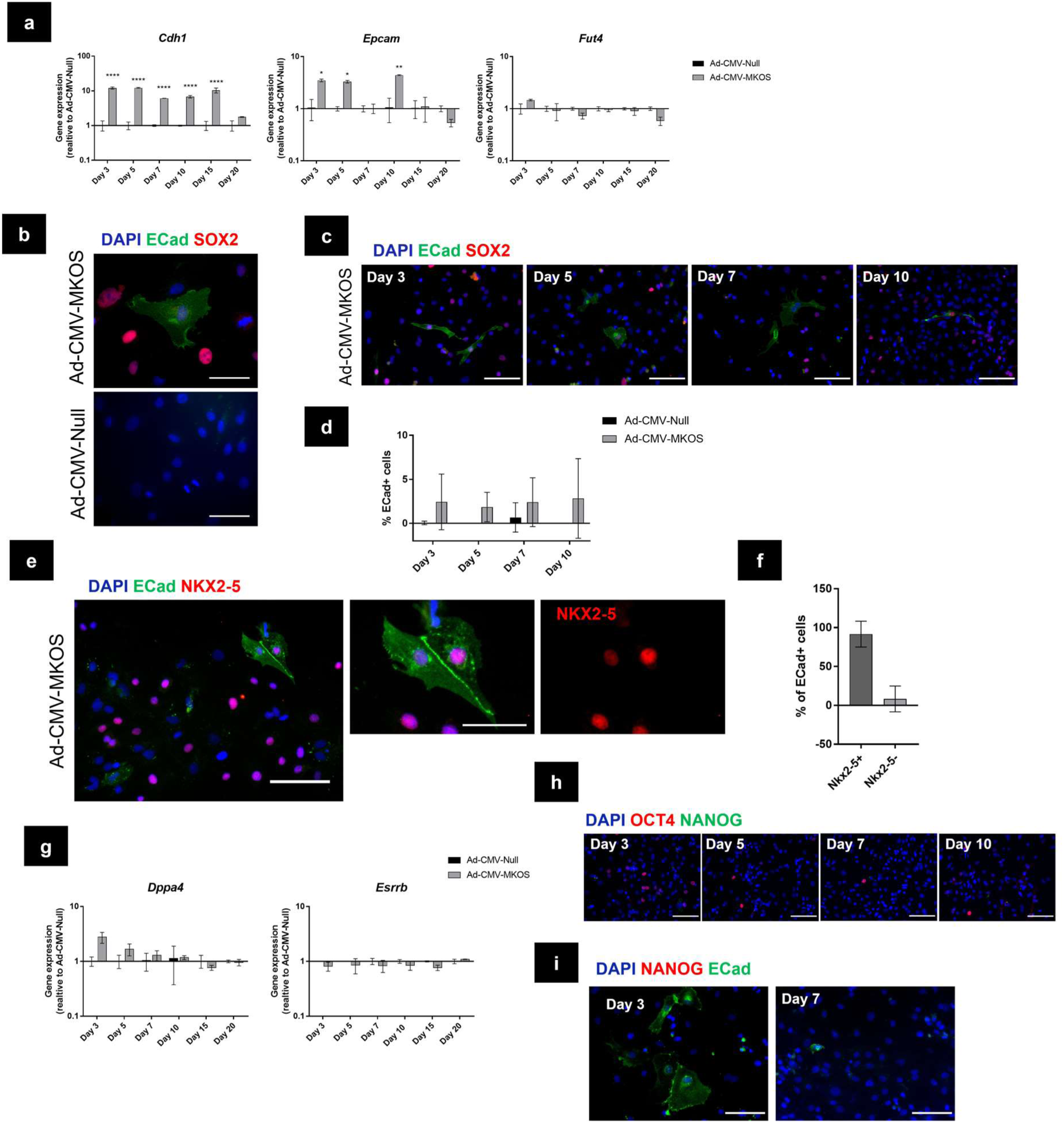
Cardiomyocytes are Partially but not Completely Reprogrammed by Adenoviral Mediated OSKM Expression. (**a**) Gene expression of genes associated with early stages of cell reprogramming to pluripotency (n=2) (**b**) Presence of ECad+ cells in Ad-CMV-MKOS treated cultures day 3 post transduction (Scale bar = 50 µm). (**c**) Ecad+ cells present at different timepoints in Ad-CMV-MKOS transduced NRCMs (Scale bar = 100 µm). (**d**) Quantification of percentage of ECad positive cells at each timepoint (n=3 replicates/3-4 fields per replicate). (**e**) Coexpression of NKX2-5 in ECad positive cells (Scale bars = 100 µm and 50 µm). (**f**) Quantification of percentage of ECad cells co-expressing NKX2-5 (n=2 replicates, 4 fields per replicate). (**g**) Expression of pluripotency genes following Ad-CMV-MKOS transduction (n=2). (**h**) Absence of NANOG expression in Ad-CMV-MKOS treated NRCMs over 10 days (scale bar = 100 µm) (n=3, 4-6 fields per replicate). (**i**) Absence of NANOG expression in ECad positive cells derived from NRCMs (scale bar = 100 µm) (n=2, 4-6 fields per replicate). Data are presented as mean ± S.D. (**a**), (**d**) and (**g**) two-way ANOVA with Tukey’s post-hoc analysis, *, **, and **** denotes p < 0.05, p < 0.01 and p < 0.0001 respectively.

We also investigated the expression of genes associated with later stages of reprogramming to pluripotency including *Esrrb* and *Dppa4* (Polo et al., 2012, Samavarchi-Tehrani et al., 2010). Both these genes are members of the core pluripotency gene regulatory network and are upregulated during maturation and stabilisation respectively (Golipour et al., 2012, Festuccia et al., 2012). Following OSKM transduction, we observed a transient increase in *Dppa4* gene expression (**Figure 4g**). However, the upregulation of this transcript was not sustained over time and we did not observe significant increases in the mRNA levels of *Esrrb* (**Figure 4g**). These results suggest that a complete pluripotency transcriptional program was not established. In addition, we did not detect NANOG protein expression at any of the time points investigated (**Figure 4h**) even when we specifically interrogated NANOG expression in cells that had initiated the MET process (**Figure 4i**). Overall, these data confirm that reprogramming of NRCMs transduced with Ad-CMV-MKOS is incomplete at least up to day 20 post transduction.

### Adenoviral Mediated Reprogramming of NRCMs is Transient

Consistent with the hypothesis that NRCMs are only partially reprogrammed by transient OSKM expression, we did not observe substantial changes in the expression of cardiogenic transcription factors *Nkx2-5, Mef2c* or *Gata4* indicating they are conserved in OSKM transduced cardiomyocytes (**Figure 5a**). However, we did observe transient decreases in the expression of *Tbx5* and cardiomyocyte contractile genes *Myh7* and *Myh6* (**Figure 5b**). The expression of these markers was at its lowest on day 3 post transduction before gradually returning toward baseline levels thereafter. In agreement, the morphological differences (i.e. dedifferentiated phenotype) observed day 3 post transduction were no longer present at the later timepoints investigated (day 15-20) when cells were indistinguishable from control vector treated NRCMs (**Figure 5c**). Furthermore, the beating capacity of the NRCM cultures was restored by day 10 post transduction and was maintained through to day 20, suggesting that partially reprogrammed cardiomyocytes regain their functionality and contractile properties (**Video S5** and **Videos S6**). Importantly, we did not detect the generation of pluripotent stem cell-like colonies at any timepoint during these investigations (**Figure 5c**). This data suggests that, in the absence of additional stimuli, partially reprogrammed cardiomyocytes have the capacity to regain a phenotype close to their original identity.

**Figure 5:**
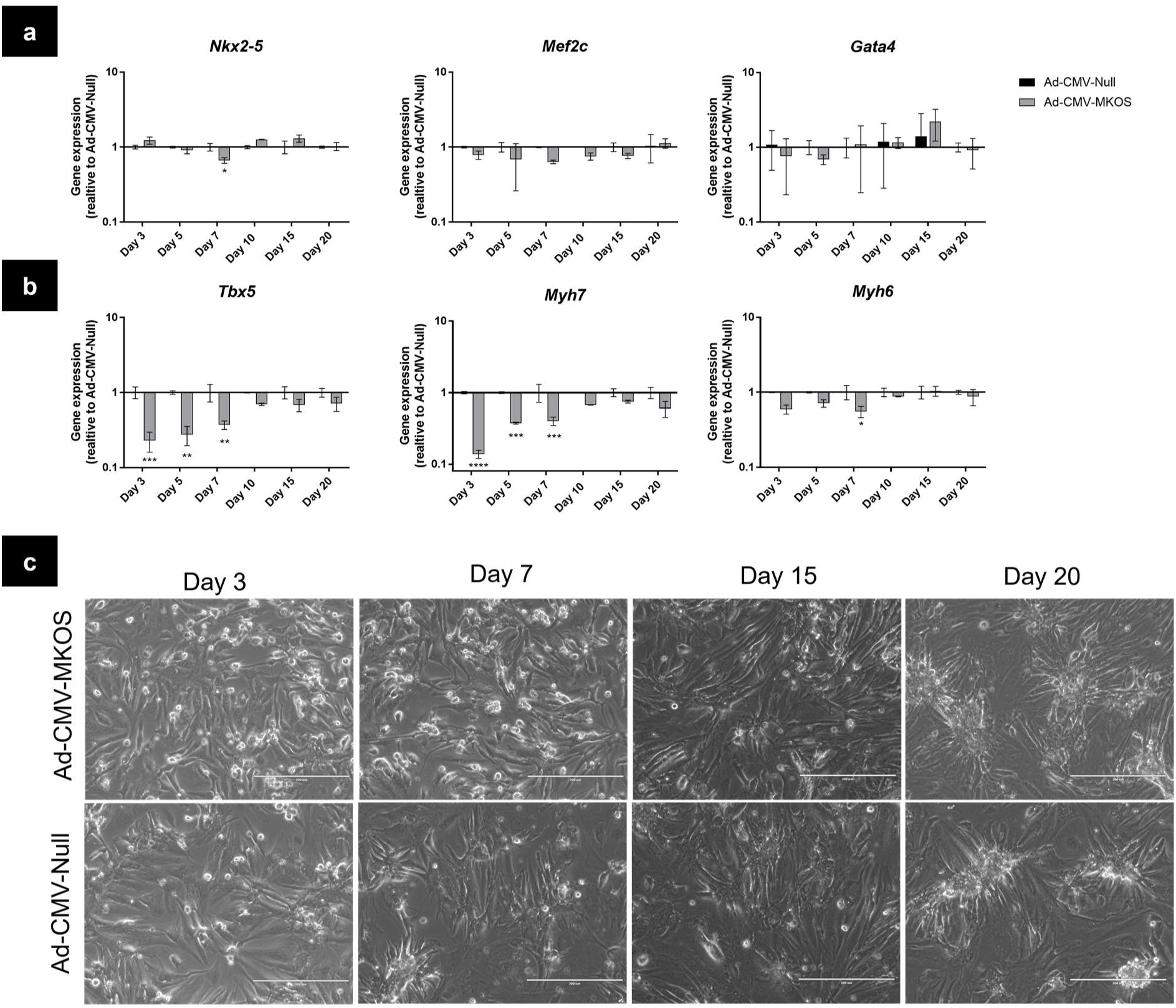
Adenoviral Mediated Reprogramming of NRCMs is Transient. (**a**) Expression of cardiogenic transcription factors after transduction with Ad-CMV-MKOS or Ad-CMV-Null (n=2). (**b**) Transient downregulation of cardiomyocyte related genes (n=2). (**c**) Phase contrast microscopy of cardiomyocytes following transduction with Ad-CMV-MKOS or Ad-CMV-Null. (scale bar = 200 µm). Representative images from n=3 replicates, 3 fields per replicate. Data are presented as mean ± S.D. (**a**) and (**b**) two-way ANOVA with Tukey’s post-hoc analysis, *, **, *** and **** denotes p < 0.05, p < 0.01, p < 0.001 and p < 0.0001 respectively.

### Partial Reprogramming of NRCMs is Enhanced in ESC Culture Media Conditions

Supplementation of cell culture medium with cytokines such as leukaemia inhibitory factor (LIF) is required for the generation and maintenance of rodent pluripotent cells (Williams et al., 1988). In addition, the replacement of FBS with chemically defined serum-free components such as KnockOut Serum Replacement (KOSR) improves iPSC generation and circumvents the variability of pro-differentiating factors present in batches of animal derived serum (Liu et al., 2014). We next investigated if culturing Ad-CMV-MKOS transduced NRCMs in ESC culture medium containing LIF and KOSR would enable the further progression of reprogramming toward pluripotency. NRCM cultures were switched to these culture conditions 2 days after viral transduction and maintained with daily media changes thereafter. As in our previous observations in cardiomyocyte (CM) media, we observed indications of dedifferentiation such as sarcomere disassembly, reduced cTnT expression, and an increase in the cTnT negative population of NRCMs (**Figure 6a**). Notably, the percentage of cardiomyocytes showing sarcomere dis-assembly was substantially greater than that previously observed in basal media indicating more efficient induction of dedifferentiation (**Figure 6b**). In agreement with this we also observed a more robust increase in the percentage of Ki67+ cells 3 days following OSKM transduction (**Figure 6c-d**). However, this difference was no longer present by day 7, suggesting that the increase in proliferation was transient (**Figure 6d**). We next compared the gene expression profiles of Ad-CMV-MKOS transduced NRCMs maintained in CM or ESC culture medium. ESC culture conditions both enhanced and prolonged the reprogramming effects triggered by OSKM as indicated by the expression of epithelial (*Cdh1, Epcam*) and cardiomyocyte (*Myh7, Tbx5, Nkx2-5*) related genes (**Figure 6e**). However, the expression of late stage reprogramming genes *Dppa4* and *Esrrb* were not increased further by this specialised media, suggesting cells were still unable to reach the later stages of reprogramming to pluripotency (**Figure 6e**). We again observed the presence of ECad+ cells however, as observed for cardiomyocyte media, the number of these cells did not appear to increase substantially over the timepoints investigated (**Figure 6f-g**). Furthermore, we were unable to detect NANOG protein expression in cells which had undergone MET (**Figure 6h**). Finally, no pluripotent stem cell-like colonies were identified for up to 20 days after Ad-CMV-MKOS transduction (**Figure S6a-b**), confirming that cell reprogramming mediated by a single transduction of a non-integrating adenoviral vector is incomplete. However, 20 days after transduction, the morphology of Ad-CMV-MKOS transduced cultures was significantly different from those treated with a control vector (**Figure S6b**). Spontaneous contractile activity was also not restored throughout the investigation indicating that in these conditions the re-differentiation of partially reprogrammed cells was limited. Together, these results indicate that media conditions are not the limiting factor on the level of reprogramming reached by NRCMs but may play a role in the efficiency and the re-differentiation of transiently reprogrammed cells. This suggests the reprogrammed state of cardiomyocytes may be maintainable in the presence of defined medias and further optimisation of culture conditions.

**Figure 6:**
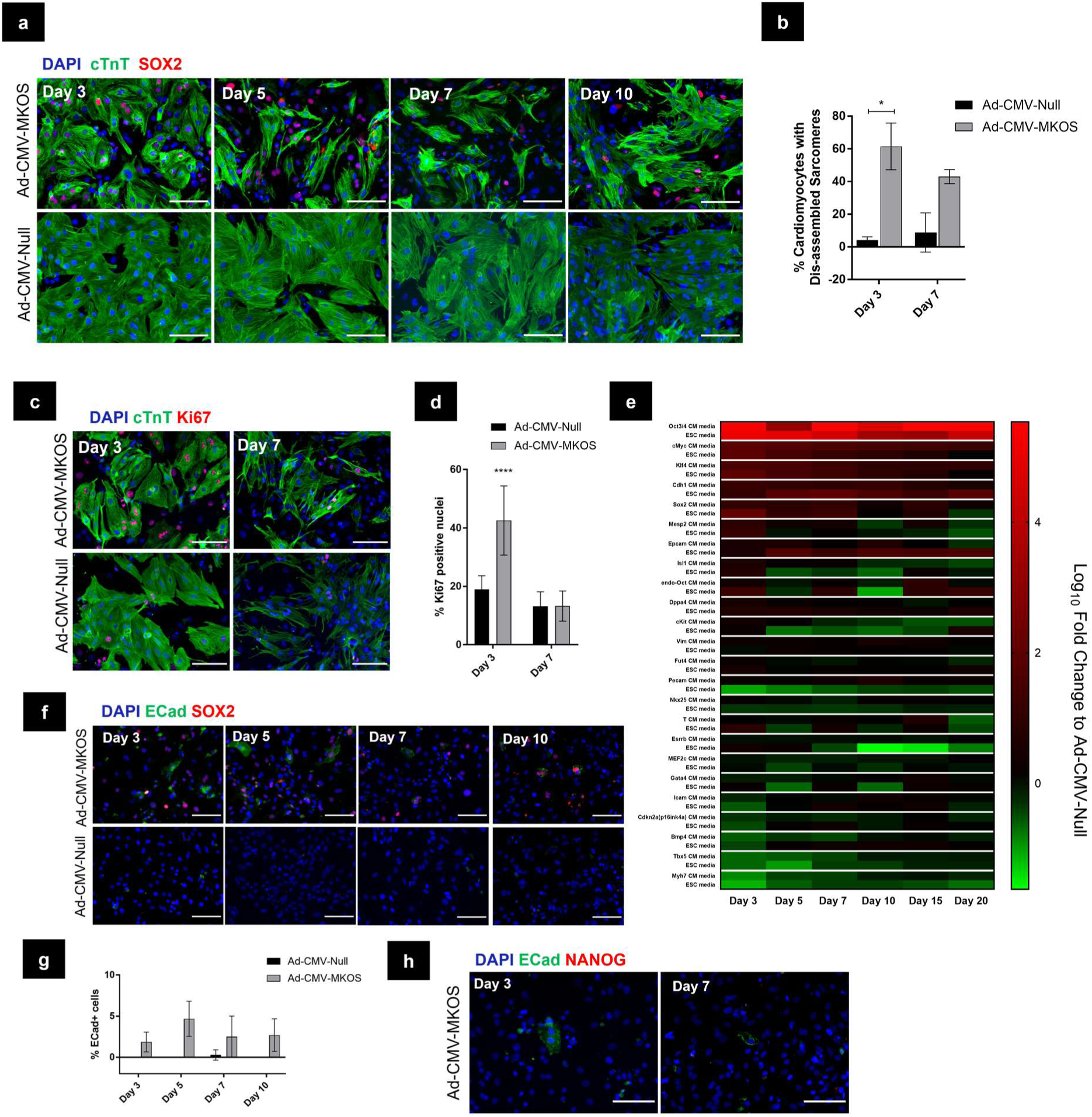
Partial Reprogramming of NRCMs is Enhanced by ESC Media Conditions. (**a**) Immunofluorescence staining of NRCMs cultured in ESC media (Scale bar = 100 µm). (**b**) Quantification of cTnT positive cells showing sarcomere dis-assembly (n=2 replicates, 4-6 fields per field). (**c**) Immunostaining of Ki67 in transduced NRCMs in ESC media (scale bar = 100 µm). (**d**) Quantification of percentage of Ki67+ cells (n=2 replicates, 4-6 fields/replicate). (**e**) RT-qPCR analysis of changes in gene expression relative to Ad-CMV-Null transduced cells in standard cardiomyocyte (CM) or ESC specified media (n=2). (**f**) Presence of ECad+ cells generated from NRCMs transduced with Ad-CMV-MKOS (Scale bar = 100 µm). (**g**) Quantification of ECad positive cells (n=2 replicates, 4-6 fields per replicate). (**h**) Absence of NANOG expression in ECad positive cells derived from NRCMs (scale bar = 100 µm). (n=2 replicates, 4-6 fields per replicate). Data are presented as mean ± S.D. (**b**), (**d**) and (**g**) two-way ANOVA with Tukey’s post-hoc analysis. * and **** denote p < 0.05 and p < 0.0001 respectively.

## Discussion

Here we demonstrate that mammalian cardiomyocytes can be partially and transiently reprogrammed to a dedifferentiated and proliferative state through adenoviral mediated OSKM expression. We chose an adenoviral vector to investigate partial cardiomyocyte reprogramming based on their established ability to efficiently transduce cardiomyocytes at low MOIs without transgene integration (Louch et al., 2011). Since proliferation is one of the early steps during cell reprogramming, we hypothesised that the OSKM transgene would be rapidly diluted in cells undergoing this process and fall below the levels required to establish and sustain pluripotency (Mikkelsen et al., 2008, Brambrink et al., 2008). Indeed, consistent with other investigations using adenoviral vectors to deliver OSKM, we did not observe the generation of pluripotent stem cells following a single adenoviral transduction (Okita et al., 2008, Zhou and Freed, 2009, Stadtfeld et al., 2008b). However, the level and extent of reprogramming achieved in cells following a single adenoviral mediated delivery of an OSKM transgene had not been previously interrogated. We confirmed the initiation of reprogramming as evidenced by cellular dedifferentiation and the induction of MET which is consistent with the responses observed following OSKM induction in other cell types (Samavarchi-Tehrani et al., 2010, Buganim et al., 2013, Tanabe et al., 2013, Mikkelsen et al., 2008). However, we did not observe NANOG+ cells arising from Ad-CMV-MKOS transduced cultures, nor the generation of pluripotent stem cell-like colonies confirming this reprogramming response to be incomplete. This is in agreement with other studies that have demonstrated the maturation phase, characterised by the establishment of pluripotency gene networks, acts as the major limiting step to the progression of reprogramming (Tanabe et al., 2013). We also observed the maintenance of NKX2-5 expression in cardiomyocytes that had initiated MET. As Nkx2-5 is one of the earliest markers of cardiogenic differentiation its persistence in this case could indicate cells are still committed to a cardiac lineage (Lien et al., 1999, Schwartz and Olson, 1999). Indeed, the expression of this gene remains relatively unchanged in the regenerating zebrafish heart where transient downregulation of other cardiac genes is observed (Raya et al., 2003, Sleep et al., 2010). Partial reprogramming of mammalian cardiomyocytes may therefore offer a method to recapitulate these limited dedifferentiation responses that contribute to the enhanced regenerative capacity of model organisms.

A single study has investigated the early stages of cardiomyocyte reprogramming to pluripotency utilising a doxycycline inducible OSKM expression system (Cheng et al., 2017). In this study the authors identified that early in the reprogramming process, prior to MET and complete loss of somatic cell identity, that genes associated with cell cycle progression are activated. Our results agree with this as both cell cycle genes were upregulated and an increase in the Ki67 positive proportion of cells was observed as early as day 3 post transduction. Furthermore, while a proportion of cells did appear to initiate MET, this was incomplete and less frequent than the number of cells undergoing dedifferentiation and proliferation. However, Cheng et al. did not interrogate the effects of early doxycycline withdrawal and instead maintained elevated OSKM expression through to the generation of stable pluripotent cells. Interestingly, in the present study, adenoviral mediated reprogramming of NRCMs was not only partial but also transient in nature. Cardiomyocyte related gene expression, sarcomere structure and striated cellular morphology was restored and cells were indistinguishable from control vector treated NRCMs by days 10-15 post transduction. This coincided with the re-establishment of autorhythmic contractile activity indicating re-differentiation of cells to a functional cardiomyocyte-like phenotype. These results are in agreement with observations in other cell types that withdrawal of exogenous OSKM expression prior to the establishment of transgene independent pluripotency enables the reversion of cells to a phenotype close to that of the starting cell population (Brambrink et al., 2008, Woltjen et al., 2009, Samavarchi-Tehrani et al., 2010, Tanabe et al., 2013, Guo et al., 2017). The presence of epigenetic memory has been identified in functionally pluripotent iPSCs that can greatly influence their differentiation potential (Kim et al., 2010, Polo et al., 2010, Rizzi et al., 2012, Xu et al., 2012, Bar-Nur et al., 2011, Shipony et al., 2014). Therefore, it is unsurprising that in cases of partial reprogramming where the epigenome is even more conserved that cells would be further restricted toward their original lineage.

Although transient OSKM mediated cardiomyocyte dedifferentiation has not been previously explored, other strategies have been investigated to enhance proliferation by inducing an immature state. This includes attempts to restore expression of genes or reactivate pathways that are inactivated during terminal differentiation of mammalian cardiomyocytes. For example forced expression of a constitutively active form of the neuregulin1 receptor ErbB2, was found induce dedifferentiation of post-natal cardiomyocytes (D’Uva et al., 2015). This ErbB2 induced dedifferentiation enhanced the proliferative capacity of both post-natal and adult cardiomyocytes such that overexpression in the heart *in vivo* enabled a more effective regenerative response. The interleukin-6 type cytokine member oncostatin M has also been shown to induce cardiomyocyte dedifferentiation and activate the re-expression of foetal and embryonic genes (Kubin et al., 2011). However, the exact mechanisms by which oncostatin M induces dedifferentiation are unclear. Inhibition of components of the Hippo pathway have also been shown to enhance cardiomyocyte proliferation by releasing the inactivation of this pathway that occurs during terminal differentiation (Xin et al., 2013, Lin et al., 2014, Leach et al., 2017, Monroe et al., 2019). A recent study demonstrated that the pro-proliferative effects of a Hippo resistant form of YAP were mediated by the reversion of cardiomyocytes to a foetal-like state by epigenetic remodelling of chromatin (Monroe et al., 2019). This epigenetic remodelling induced by YAP indicate parallels between these endogenous dedifferentiation mechanisms and with that of reprogramming induced by OSKM. Overall, these studies in combination with the findings presented here further support the hypothesis that inducing dedifferentiation of cardiomyocytes to more immature states can enhance the proliferative potential of these cells. Transient OSKM mediated reprogramming therefore offers a novel tool for further exploration of induced dedifferentiation in cardiac regenerative medicine.

Given the similarities between OSKM induced transient reprogramming presented here and the endogenous regenerative mechanisms observed in model organisms (Poss et al., 2002, Jopling et al., 2010, Sleep et al., 2010), transient OSKM expression could be a novel strategy to stimulate proliferation of cells in the injured myocardium *in vivo*. Some of us have previously demonstrated that somatic cells in liver and skeletal muscle can be transiently reprogrammed *in vivo* through non-integrating vector mediated expression of OSKM (Yilmazer et al., 2013a, Yilmazer et al., 2013b, de Lázaro et al., 2014, de Lázaro et al., 2019). Consistent with the results observed by others, transiently reprogrammed cells appear to undergo a short proliferative stage before undergoing re-differentiation and integration into the host tissue (de Lázaro et al., 2019, Ohnishi et al., 2014). Ourselves and others have also demonstrated that transient *in vivo* reprogramming can be used to enhance the regeneration of injured tissue (Seo et al., 2016, de Lázaro et al., 2019, Doeser et al., 2018) and rejuvenate aged tissues (Ocampo et al., 2016).

The present study demonstrates for the first time that transient OSKM expression in cardiomyocytes *in vitro* enables a temporary passage through a proliferative de-differentiated state which is encouraging for the potential applications of this approach in the heart. Furthermore, in contrast to investigations relying on inducible transgenic models (Seo et al., 2016, Ocampo et al., 2016), the use of an adenoviral vector to induce transient reprogramming offers a more readily translatable approach for *in vivo* applications.

We have also identified several limitations in the present study that would be important to address in future investigations. It will be important to confirm that adenoviral mediated transient reprogramming can be induced in adult cardiomyocytes and can overcome the more complete exit from the cell cycle that occurs in these cells. Further to this, although we observed indications that a proportion of reprogrammed cardiomyocytes reverted to functional cardiomyocyte-like cells we cannot confirm the fate of all cells transduced with Ad-CMV-MKOS. It will therefore be important that future investigations utilise more complete lineage tracing approaches to confirm the fate of partially reprogrammed cells and determine the level of heterogeneity in the final cell population. Finally, the enhancement of cardiomyocyte de-differentiation in ESC media and the prevention of re-differentiation to a beating cardiomyocyte-like state suggests further optimization of media conditions could be utilised to prolong the de-differentiated state of partially reprogrammed cardiomyocytes. This could offer a novel tool to generate expandable progenitor-like populations for both basic and translation research applications.

## Conclusion

This study offers previously unreported demonstration of adenoviral-mediated OSKM expression to enable transient dedifferentiation and proliferation of primary cardiomyocytes. This, in many ways, recapitulates what is seen during zebrafish and neonatal mammalian regeneration. We therefore propose that it could be utilised further as an investigative tool to inform cardiac regenerative strategies that aim to enhance the proliferation of cardiomyocytes *in situ*.

## Experimental

### Animals

All animal experiments were performed with prior approval from the UK Home Office under the project license (P089E2EOA) and in compliance with the UK Animals (Scientific Procedures) Act, 1986. Time mated Sprague-Dawley rats were obtained from Charles River (UK) and allowed at least 1 week to acclimatise before litters were born. αMHC-Cre-tdTomato (C57BL6/129/FVB) mice were derived from crossing αMHC-Cre (C57BL6/129/FVB) mice, kindly provided by Dr Elizabeth Cartwright (University of Manchester, UK) with B6.Cg-Gt(ROSA)26Sortm14(CAG-tdTomato)Hze/J. The resultant mouse strain αMHC-Cre-tdTomato (C57BL6/129/FVB) was maintained as heterozygous for the αMHC-Cre transgene gene and homozygous for ROSA26tdTomato.

### Viral vectors

The viral vectors used in this study were purchased from Vector Biolabs (USA). Ad-CMV-MKOS is a Human Adenovirus type 5 (dE1/E3) containing the polycistronic expression cassette encoding mouse c-Myc-F2A-Klf4-T2A-Oct4-E2A-Sox2 under the control of a CMV promoter. A serotype matched adenovirus containing an empty CMV promoter expression cassette was used as a control. Vectors were provided in phosphate buffered saline (PBS) containing 5% glycerol and were diluted in culture media immediately prior to transduction.

### Primary neonatal rat cardiomyocyte (NRCM) extraction

NRCMs were extracted from 2 day old Sprague-Dawley (SD) rats (Charles River, UK) as described previously with some modifications (Mohamed et al., 2016). Briefly, animals were culled by cervical dislocation followed by decapitation, hearts were excised and washed twice in ADS buffer (116 mM NaCl, 20 mM HEPES, 1 mM NaH_2_PO_4_, 5.5 mM glucose, 5.5 mM KCl and 1 mM MgSO_4_, pH 7.4). Ventricles were dissected from the atria and cut laterally. Next, they were digested in 7 ml ADS buffer containing 0.6 mg/ml collagenase A (Roche, Germany) and 0.6 mg/ml pancreatin (Sigma-Aldrich, UK) at 37 °C for 5 minutes, under moderate shaking. Digested tissue was then passed several times through a 25 ml glass pipette with the resulting supernatant collected and passed through a 70 µm cell strainer (Corning, UK). Enzymatic digestion was inactivated with 3 ml fetal bovine serum (FBS) (ThermoFisher, UK). This process was repeated 9 times or until ventricular tissue had been digested completely. The resulting cell solution was centrifuged at 1200 rpm, re-suspended in pre-plating medium (68% Dulbecco’s Modified Eagle Medium (DMEM, Sigma-Aldrich, UK), 17% Medium-199 (M199, Sigma-Aldrich, UK) supplemented with 10% horse serum (ThermoFisher, UK), 5% FBS and 2.5 μg/ml amphotericin B (Sigma-Aldrich, UK)) and plated on 90 mm tissue culture dishes (Corning, UK). Non-myocytes were left to adhere for 1 hour and the cardiomyocyte enriched supernatant was then collected. The resulting NRCMs were counted and plated at 0.2×10^6^ cells/well in Corning Primaria 24 well plates, 8×10^4^ cells/well in poly-l-lysine (Sigma-Aldrich, UK)/Laminin (Sigma-Aldrich, UK) coated MilliCell EZ 8 well chambers slides (Merck, UK) and 3×10^4^ cells/well in Corning Primaria 96 well plates. NRCMs were initially maintained in pre-plating medium with the addition of 1 µM BrdU (Sigma-Aldrich, UK) at 37 °C for 24 hours. Cells were then washed twice in warm PBS containing Ca^2+^ and Mg^2+^ (Gibco, UK) and medium replaced with cardiomyocyte maintenance medium (80% DMEM and 20% Medium 199 supplemented with 1% FBS, 2.5 μg/ml amphotericin B and 1 μM BrdU) in which cells were maintained until viral transduction.

### Adenoviral transduction and culturing

NRCMs were transduced with Ad-CMV-MKOS (Vector Biolabs) or Ad-CMV-Null (Vector Biolabs) at MOIs of 1-10 PFU/cell pre-diluted in cardiomyocyte maintenance media containing 2% FBS without BrdU. NRCMs were then maintained at 37 °C for 24 hours before virus containing media was replaced with fresh cardiomyocyte maintenance media (without BrdU). Media was exchanged every other day thereafter. In a subset of experiments, as indicated in the text, media was replaced with ESC medium (Knockout DMEM/F12 (ThermoFisher, UK) supplemented with 15% Knockout Serum Replacement (ThermoFisher, UK), 1% StemPro non-essential amino acids (ThermoFisher, UK), 50 µM β-mercaptoethanol (Sigma-Aldrich, UK), 10 ng/ml of mouse leukaemia inhibitory factor (eBioscience, UK) and was changed daily thereafter.

### αMHC-Cre-tdTomato NMCMs

NMCMs were extracted from post-natal day 2 pups using a modified version of the protocol for NRCMs. In brief, prior to tissue digestion and cell extraction, hearts were screened with an EVOS FL fluorescence microscope to select hearts that were positive for tdTomato fluorescence confirming a positive αMHC-Cre genotype. Following extraction, NMCMs were counted and seeded at 8×10^4^ cells/well in poly-l-lysine/laminin coated MilliCell EZ 8 well chambers slides. Transduction and maintenance of NMCMs was performed as described above for NRCMs.

### Imaging/Videos

Still phase contrast images were taken on an EVOS FL microscope using the 10x and 20x objective. Videos were captured on an Olympus IX83 inverted microscope using a 20x objective and captured using an Orca ER camera (Hamamatsu, UK) through MMI Cell tools software (MMI, Switzerland).

### alamarBlue assay and cell counts

Resazurin sodium salt (Sigma-Aldrich, UK) was dissolved in PBS containing Ca^2+^ and Mg^2+^ which was further diluted in cell culture medium to a final concentration of 20 µg/ml, immediately before adding to the cells in 96 well plates. Cells were then incubated for 2 hours at 37 °C in the absence of light. Fluorescence was read directly from plates using a FLUOstar Omega (BMG Labtech, UK) plate reader with 544 nm excitation and 590 nm emission wavelengths. The percentage cell viability relative to untreated cells was calculated using the mean fluorescence intensity measurement from 6 replicates per condition. Following fluorescence measurements cells were washed in PBS, detached using trypsin-EDTA and counted using trypan blue exclusion with a haematocytometer.

### Reverse-transcription real-time quantitative polymerase chain reaction (RT-qPCR)

Cultured cells were lysed directly in tissue culture plates in lysis buffer (ThermoFisher, UK) containing 1% β-mercaptoethanol. RNA was then extracted from the cells or tissues using the PureLink RNA mini kit (ThermoFisher, UK) following the manufacturer’s guidelines. Eluted RNA was subjected to an additional DNase treatment using the RapidOut DNA Removal kit (ThermoFisher, UK) to remove any potentially contaminating viral DNA. Purified RNA was quantified using a biophotometer and 0.8-1 µg of RNA was used to produce cDNA using the High Capacity Reverse Transcription kit (ThermoFisher, UK) following manufacturer’s protocol. cDNA samples (2 µl) were combined with primers and PowerUp SYBR Green Mastermix (ThermoFisher, UK) following manufacturer’s instructions. RT-qPCR reactions were run in duplicate on a BioRad CFX thermal cycler (BioRad, UK) according to the following protocol: 50 °C for 2 minutes, 95 °C for 2 minutes and 40 cycles of 95 °C 15 seconds and 60 °C for 1 minute. Melt curve analysis was included to ensure amplification of a single PCR product and non-reverse transcribed controls were used to confirm no contamination with viral or genomic DNA. Data was analysed using the Livak method (2^-ΔΔCt^) using *β-actin* as a housekeeping gene and normalising to the relevant controls for each experiment (Livak and Schmittgen, 2001). High throughput dynamic array RT-qPCR was also carried out using a Biomark HD (Fluidigm, UK) in the 96.96 format. In this case data was analysed using the Livak method (2^-ΔΔCt^) using *Gapdh* as a housekeeping gene and normalising to the Ad-CMV-Null treated cells. Primer pairs used are provided in **Supplementary Table 1**.

### Immunocytochemistry

Cells grown on poly-l-lysine/laminin coated 8 well MilliCell EZ slides were fixed in ice cold 4% PFA (Sigma-Aldrich, UK) for 15 minutes, permeabilized with PBS containing 0.3% Triton-X (Sigma-Aldrich, UK) for 10 minutes and blocked in PBS containing 0.1% Tween20 (PBST) with 10% normal goat serum (NGS) (ThermoFisher, UK) and 0.3M glycine (Sigma-Aldrich, UK) for 1 hour. Cells were then incubated with primary antibodies in PBST containing 10% NGS overnight at 4 °C in a humidified chamber. The following day, cells were washed in PBST and incubated in secondary antibody diluted in PBST containing 10% NGS for 1 hour at room temperature. Cells were washed with 3 times with PBST, twice with PBS and mounted with ProLong Gold mounting reagent with DAPI (ThermoFisher, UK) which was allowed to cure for 24 hours prior to imaging. Fluorescent slides were imaged on a Zeiss AXIO Observser. A1 using a 10x or 20x objective. Higher magnification images were acquired using an Olympus IX83 inverted microscope using the 60x objective with a Z optical spacing of 0.2 μm. Images were analysed and quantified using ImageJ (NIH, USA). Where appropriate, brightness/contrast for individual fluorescence channels was adjusted equally between experimental samples and controls. The antibodies used in these investigations are provided in **Supplementary Table 2**.

### Statistical analysis

Statistical analysis was carried out using GraphPad Prism 8 to perform unpaired t-test’s for comparisons between 2 groups with correction for multiple comparisons where appropriate, or ANOVA followed by Tukey’s post-hoc analysis for comparison between 3 or more groups. A probability (P) of < 0.05 was regarded as statistically significant and P values and n numbers are specified in the figure legends.

## Acknowledgements

T.K would like to thank the EPSRC & MRC CDT in Regenerative Medicine (EP/L014904/1) for a funded PhD studentship. T.K, I.d.L and K.K. would also like to acknowledge the generous gift to the Nanomedicine Lab towards development of this project by an anonymous donor. The authors would also like to acknowledge the Genomic technologies and Bioimaging core facilities at the University of Manchester, UK.

## Author contributions

T.K, I.d.L and K.K. conceived the project and designed experiments. T.K. performed the majority of experiments. M.S. performed some immunofluorescence experiments. G.C. advised in experimental design and interpretation of data. T.K., I.d.L, G.C. and K.K. wrote the manuscript.

## Supplementary Information

### Supplementary Figures

**Figure S1:** (**a**) Representative phase contrast and immunofluorescence images of NRCMs on day 0 of transduction (Scale bar = 200 µm). (**b**) Quantification of cardiomyocytes (cTnT+/VIM-) and non-myocytes (VIM+/cTnT-) as a percentage of total population (n=8 fields). (**c**) Co-expression of NKX2-5 exclusively in cTnT positive cells (Scale bar = 200 µm). (b) Data presented as mean percentage ± S.D.

**Figure S2**: (**a**) Gene expression of *Myh6* and *Myh7* in NRCMs treated with Ad-CMV-Null (n=3). Data are presented as mean ± S.D. one-way ANOVA with Tukey’s post hoc analysis. No statistically significant differences were observed.

**Figure S3**: (**a**) Immunofluorescence of αMHC-Cre-tdTomato NMCMs 3 days post transduction with either Ad-CMV-Null or Ad-CMV-MKOS showing the presence of dedifferentiating cardiomyocytes (yellow arrows) (scale bars = 50 µm). Representative image from n=2 replicates, 4-6 fields per replicate.

**Figure S4:** (**a**) Ki67 expression in αMHC-Cre-tdTomato cardiomyocytes 3 days post transduction (Scale bar = 100 µm). (**b**) Quantification of Ki67+ nuclei and tdTomato+ Ki67+ cells (n=2 replicates/4 fields per replicate). (**c**) Expression of Ki67 in cTnT-tdTomato+ cells 3 days post transduction (Scale bar = 50 µm). Data are presented as mean ± S.D. (**b**) Unpaired t-tests, no statistically significant differences identified.

**Figure S5:** (**a**) Presence of ECad+ tdTomato+ cells in αMHC-Cre-tdTomato cardiomyocytes 5 days post transduction (Scale bar = 100 µm). (**b**) High magnification of ECad+ cells with orthogonal view to confirm co-localisation with tdTomato (Scale bar = 50 µm).

**Figure S6:** (**a**) Phase contrast microscopy of NRCMs days 3-15 post transduction with Ad-CMV-MKOS in the presence of ESC media (Scale bar = 200 µm). (**b**) Lack of ESC-like colonies in NRCMs transduced with Ad-CMV-MKOS day 20 post transduction (Scale bar = 400 µm). Representative images from n=3 repeats/4 fields per repeat.

### Supplementary Videos

**Video S1:** Ad-CMV-Null treated cardiomyocytes day 3 post transduction.

**Video S2:** Ad-CMV-MKOS treated cardiomyocytes day 3 post transduction.

**Video S3:** Ad-CMV-Null treated cardiomyocytes day 10 post transduction.

**Video S4:** Ad-CMV-MKOS treated cardiomyocytes day 10 post transduction.

**Video S5:** Ad-CMV-Null treated cardiomyocytes day 20 post transduction.

**Video S6:** Ad-CMV-MKOS treated cardiomyocytes day 20 post transduction.

### Supplementary Tables

**Table S1:** Primer pairs utilised in RT-qPCR investigations.

**Table S2:** Antibodies utilised in immunocytochemistry investigations.

