## Supplementary Information for "Transient Reprogramming of Neonatal Cardiomyocytes to a Proliferative Dedifferentiated State"

### FIGURE S1

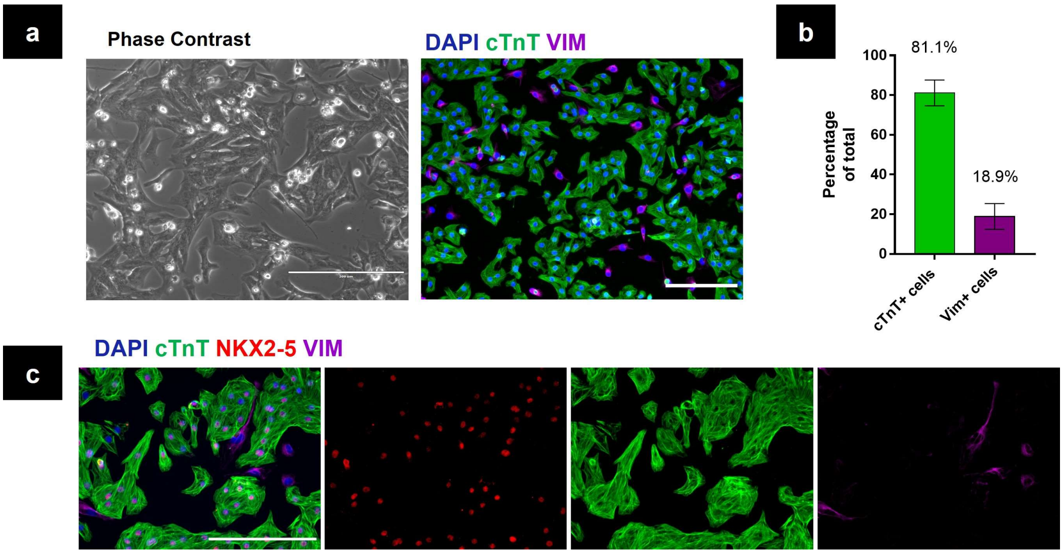


**Figure S1:** (**a**) Representative phase contrast and immunofluorescence images of NRCMs on day 0 of transduction (Scale bar = 200 µm). (**b**) Quantification of cardiomyocytes (cTnT+/VIM-) and non-myocytes (VIM+/cTnT-) as a percentage of total population (n=8 fields). (**c**) Co-expression of NKX2-5 exclusively in cTnT positive cells (Scale bar = 200 µm).
(b) Data presented as mean percentage ± S.D.

### FIGURE S2

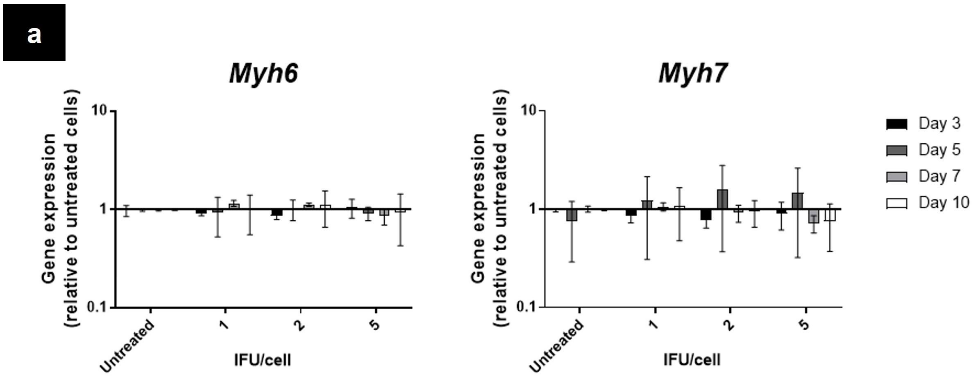


**Figure S2**: (**a**) Gene expression of *Myh6* and *Myh7* in NRCMs treated with Ad-CMV-Null (n=3).
Data are presented as mean ± S.D. one-way ANOVA with Tukey's post hoc analysis. No statistically significant differences were observed.

### FIGURE S3

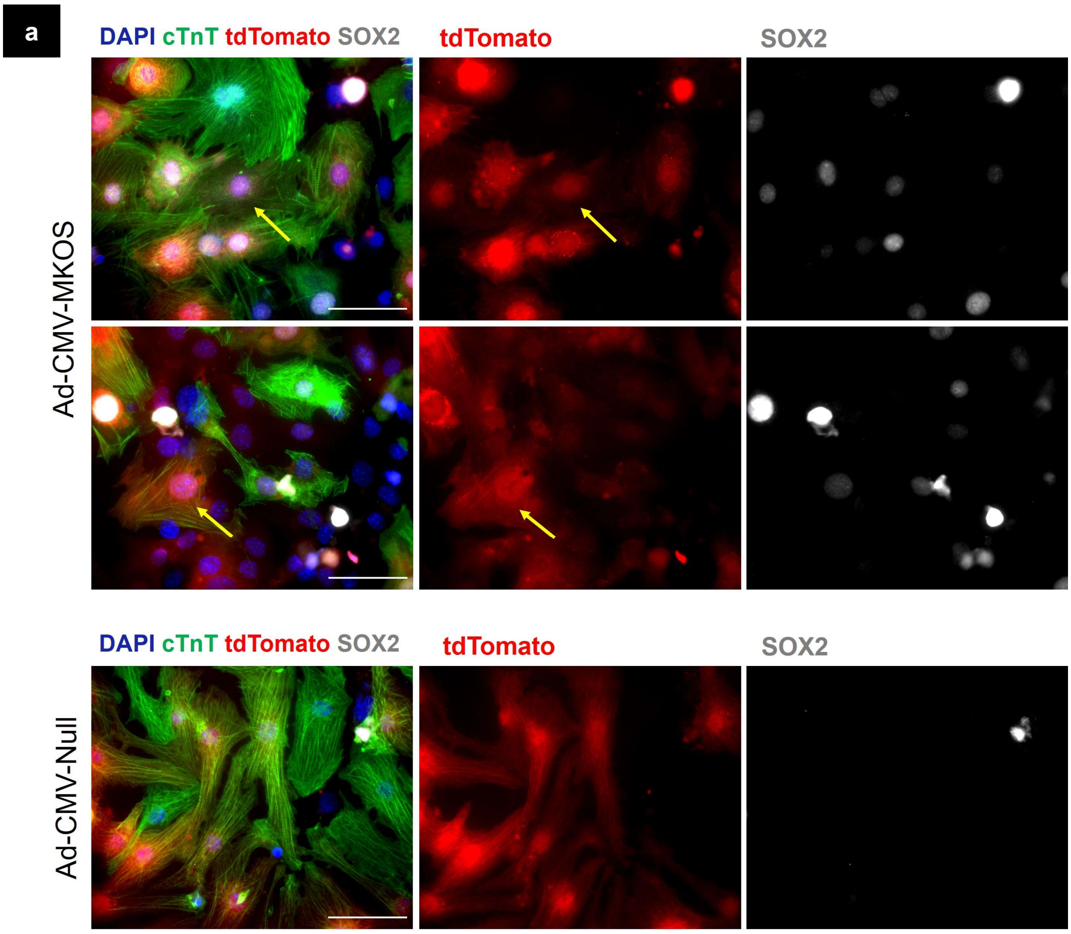


**Figure S3**: (**a**) Immunofluorescence of αMHC-Cre-tdTomato NMCMs 3 days post transduction with either Ad-CMV-Null or Ad-CMV-MKOS showing the presence of dedifferentiating cardiomyocytes (yellow arrows) (scale bars = 50 μm). Representative image from n=2 replicates, 4-6 fields per replicate.

### FIGURE S4

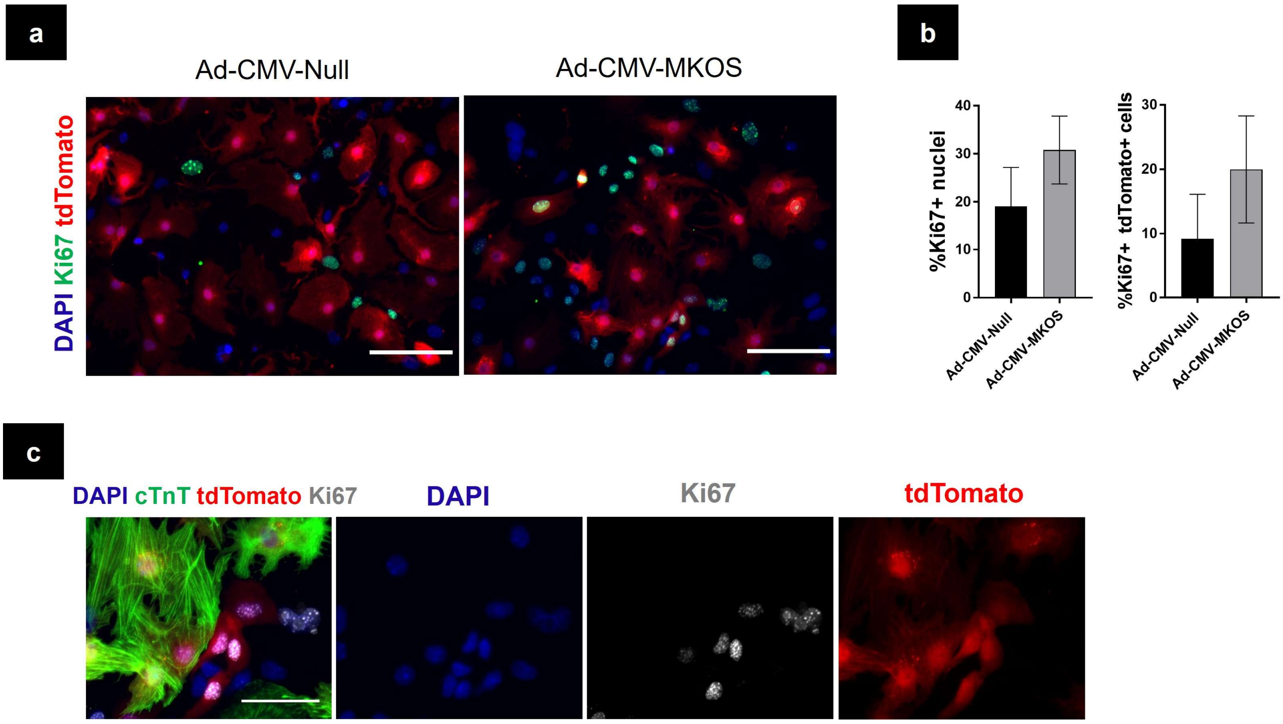


**Figure S4:** (**a**) Ki67 expression in αMHC-Cre-tdTomato cardiomyocytes 3 days post transduction (Scale bar = 100 μm). (**b**) Quantification of Ki67+ nuclei and tdTomato+ Ki67+ cells (n=2 replicates/4 fields per replicate). (**c**) Expression of Ki67 in cTnT- tdTomato+ cells 3 days post transduction (Scale bar = 50 μm).
Data are presented as mean ± S.D. (**b**) Unpaired t-tests, no statistically significant differences identified.

### FIGURE S5

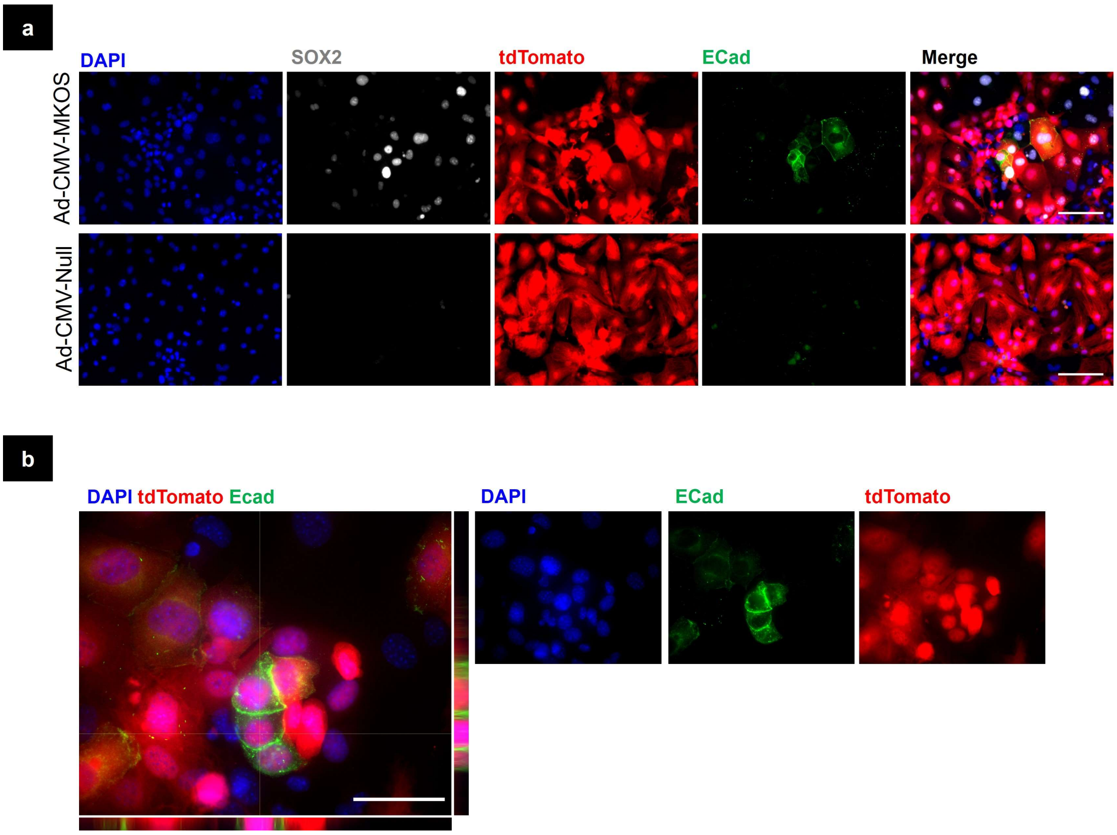


**Figure S5:** (**a**) Presence of ECad+ tdTomato+ cells in αMHC-Cre-tdTomato cardiomyocytes 5 days post transduction (Scale bar = 100 µm). (**b**) High magnification of ECad+ cells with orthogonal view to confirm co-localisation with tdTomato (Scale bar = 50 µm).

FIGURE S6

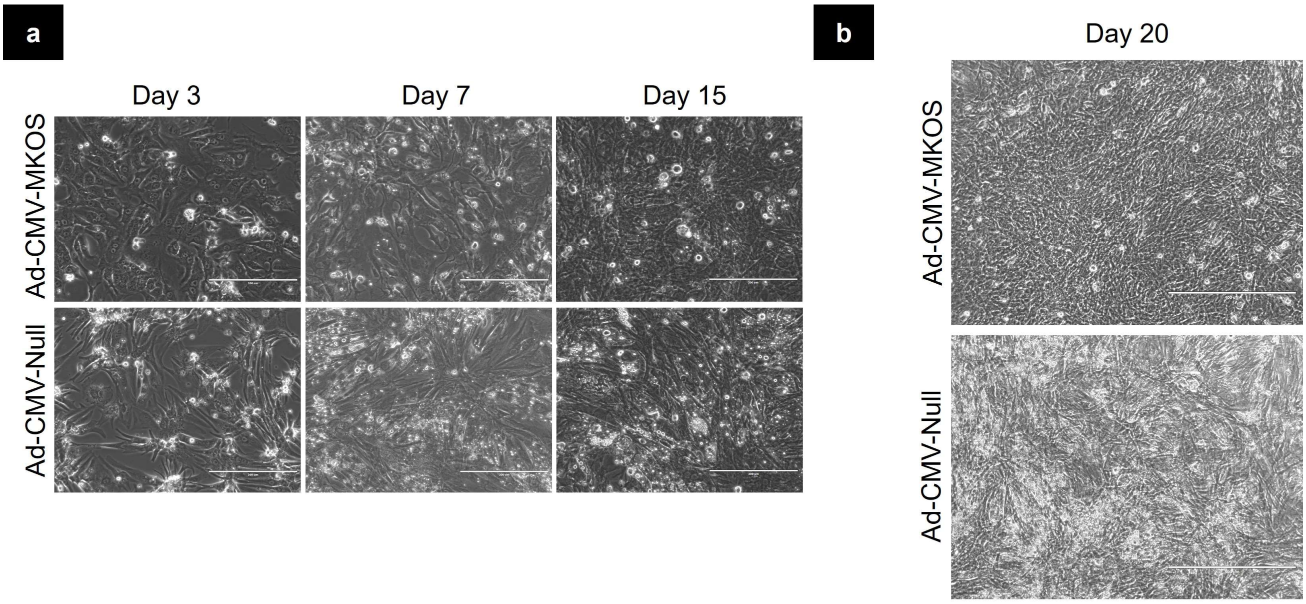


**Figure S6:** (**a**) Phase contrast microscopy of NRCMs days 3-15 post transduction with Ad-CMV-MKOS in the presence of ESC media (Scale bar = 200 µm). (**b**) Lack of ESC-like colonies in NRCMs transduced with Ad-CMV-MKOS day 20 post transduction (Scale bar = 400 µm). Representative images from n=3 repeats/4 fields per repeat.

#### Supplementary Videos

**Video S1:** Ad-CMV-Null treated cardiomyocytes day 3 post transduction

**Video S2:** Ad-CMV-MKOS treated cardiomyocytes day 3 post transduction

**Video S3:** Ad-CMV-Null treated cardiomyocytes day 10 post transduction

**Video S4:** Ad-CMV-MKOS treated cardiomyocytes day 10 post transduction

**Video S5:** Ad-CMV-Null treated cardiomyocytes day 20 post transduction

**Video S6:** Ad-CMV-MKOS treated cardiomyocytes day 20 post transduction

#### Supplementary Tables

**Table S1:** Primer pairs utilised in RT-qPCR investigations.

**Table S2:** Antibodies utilised in immunocytochemistry investigations.
