## Supplementary Tables for "Transient Reprogramming of Neonatal Cardiomyocytes to a Proliferative Dedifferentiated State"

| *Gene symbol* | **Forward Primer (5'->3')** | **Reverse Primer (5'->3')** |
|---|---|---|
| *Oct3/4 (total)* | TGAGAACCTTCAGGAGATATGCAA | CTCAATGCTAGTTCGCTTTCTCTTC |
| *Oct3/4 (endogenous)* | TCTTTCCACCAGGCCCCCGGCTC | TGCGGGCGGACATGGGGAGATCC |
| *Sox2 (total)* | GGTTACCTCTTCCTCCCACTCCAG | TCACATGTGCGACAGGGGCAG |
| *Klf4 (total)* | CAGTGGTAAGGTTTCTCGCC | GCCACCCACACTTGTGACTA |
| *cMyc (total)* | CAGAGGAGGAACGAGCTGAAGCGC | TTATGCACCAGAGTTTCGAAGCTGTTCG |
| *Actb* | GACCTCTATGCCAACACAGT | AGTACTTGCGCTCAGGAGGA |
| *Alpl* | ATCGGACCCTGCCTTACCAA | GGAGACGCCCATACCATCTC |
| *Bmp4* | TGGACACCTCATCACACGACTA | GCGACGGCAGTTCTTATTCTTC |
| *Ccna2* | AGTGCCGCTGTCTCTTTACC | GGGGTGATTCAAAACTACCATCC |
| *Ccnd1* | TCAAGTGTGACCCGGACTG | GACCAGCTTCTTCCTCCACTT |
| *Cdh1* | TTCAACCCAAGCACGTACCA | CAGAATGCCCTCGTTGGTCT |
| *Cdkn1a* | CACGGCTCAGTGGACCAGAA | ACTGGAGCTGCCTGAGGTAGGA |
| *Cdkn2a* | CGTACCCCGATACAGGTGATG | ACCAGAGTGTCTAGGAAGCCC |
| *Dppa4* | CTTTCCCAGATCAAATGCCCG | TCTTCCCCAGGTTCCGTCTT |
| *Epcam* | CGGGGATTGTTGTCCTGGTT | CCCATCTCCTTTATCTCAGCCTTC |
| *Esrrb* | GGGATGCTGAAGGAAGGCGT | TTAGCAGGCGGGGAAATCTG |
| *Fut4* | GCTGTCTATCGCCGCTACTT | TGCCAAGTTGTGGATGCTCT |
| *Gapdh* | ACCACAGTCCATGCCATCAC | TCCACCACCCTGTTGCTGTA |
| *Gata4* | CTCTATCACAAGATGAACGGCATCAA | TCTGGCAGTTGGCACAGGAGAG |
| *Isl1* | CTGCAAATGGCAGCCGAGC | GGTCTTCTCGGGCTGTTTGT |
| *Kit* | CTCCAACGATGTGGGCAAGA | GGGCCTGGATTTGCTCTTTG |
| *Mef2c* | AGGCACCAGCGCAGGGAATG | CCACCGGGGTAGCCAATGACT |
| *Mesp1* | CATTTAAGCCCGGTTGCCTG | TGCTGAAGAGCGGAGACGAG |
| *Mesp2* | AACAAGACTGGGCACTGGAC | CTGGAGACACAGAAAGACTCTGG |
| *Myh6* | TAACCGGAGTTTAAGAGTGACAGG | TAGGCGCTCCTTCTCTGACT |
| *Myh7* | CTGGCACCGTGGACTACAAT | GCCCTTGTCTACAGGTGCAT |
| *Nkx2-5* | ACCGCCCCTACATTTTATCCG | CACAGCTCTTTCTTATCCGCCC |
| *Pecam1* | CTGCCAGTCAGTAAATGGGAC | CTTCATCCACCGGGGCTATT |
| *Sox17* | GGCACGGAACCCAACCAGC | CAGTCGTGTCCCTGGTAGGGAAGAC |
| *Tbx5* | GGTCCGTAACTGGTAAAG | ATTTTCGTCTGCTTTCAC |
| *Tbxt (T)* | AGAATGAGGAGATTACGGCCC | ATTGGGAATATCCCGGCTGC |
| *Vim* | GCGAGAGAAATTGCAGGAGGA | CGTTCAAGGTCAAGACGTGC |

**Table S1**: Primer pairs used for RT-qPCR investigations.

| Primary antibodies | | | |
|---|---|---|---|
| **Antibody** | **Species** | **Supplier** (Catalogue number) | **Dilution used** |
| Anti-Sox2 | Rabbit | Abcam (ab97959) | 1:500 |
| Anti-Oct3/4 | Rabbit | Abcam (ab19857) | 1:250 |
| Anti-Oct3/4 | Rat | ThermoFisher (14-5841-82) | 1:250 |
| Anti-Nanog | Rabbit | Abcam (ab80892) | 1:200 |
| Anti-Nanog | Rabbit | Abcam (ab106465) | 1:200 |
| Anti-Cardiac TroponinT | Mouse | Abcam (ab8295) | 1:500 |
| Anti-Nkx2-5 | Rabbit | ProteinTech (13921-1-AP) | 1:300 |
| Anti-ECadherin | Mouse | Abcam (ab76055) | 1:200 |
| Anti-Ki67 | Rabbit | Abcam (ab15580) | 1:800 |
| Anti-Vimentin | Chicken | Abcam (ab24525) | 1:1000 |
| **Secondary antibodies** | | | |
| **Antibody** | **Species** | **Supplier (Catalogue number)** | **Dilution used** |
| Anti-Mouse Alexa Fluor 488 | Goat | ThermoFisher (A11001) | 1:500 |
| Anti-Rabbit Alexa Fluor 594 | Goat | ThermoFisher (A11012) | 1:500 |
| Anti-Rat Alexa Fluor 647 | Goat | ThermoFisher (A21247) | 1:300 |
| Anti-Chicken Alexa Fluor 647 | Goat | ThermoFisher (A21449) | 1:1000 |

**Table S2:** Antibodies used for immunocytochemistry investigations.
